# Immune determinants of the association between tumor mutational burden and immunotherapy response across cancer types

**DOI:** 10.1101/2021.05.25.445197

**Authors:** Neelam Sinha, Sanju Sinha, Cristina Valero, Alejandro A. Schäffer, Kenneth Aldape, Kevin Litchfield, Timothy A Chan, Luc G T Morris, Eytan Ruppin

## Abstract

The FDA has recently approved high tumor mutational burden (TMB), defined by ≥10 mutations/Mb, as a biomarker for the treatment of solid tumors with pembrolizumab, an immune checkpoint inhibitor (ICI) that targets PD1. However, recent studies testify that high TMB levels are only able to stratify ICI responders in a subset of cancer types, where the mechanisms underlying this observation have remained unknown. We hypothesized that the tumor immune microenvironment (TME) may modulate the stratification power of TMB (termed <u>*TMB power*</u>) in a cancer type, leading to this observation. To systematically study this hypothesis, we analyzed TCGA expression data to infer the levels of 31 immune-related factors characteristic of the TME of different cancer types. We integrated this information with TMB and response data of 2,277 patients treated with anti-PD1 or anti-PD-L1 ICI to identify the key immune factors that can determine *TMB power* across 14 different cancer types. We find that high levels of M1 macrophages and low resting dendritic cells in the TME characterize cancer types with high *TMB power.* A model based on these two immune factors is strongly predictive of the *TMB power* in a given cancer type (Spearman Rho=0.76, P<3.6×10^−04^). Using this model, we provide predictions of the *TMB power* in nine additional cancer types, including rare cancers, for which TMB and ICI response data are not yet publicly available on a large scale. Our analysis indicates that TMB-High may be highly predictive of ICI response in cervical squamous cell carcinoma, suggesting that such a study should be prioritized.

## Main

Immunotherapy has shown remarkable clinical benefit in many cancers. However, its benefit is limited to a subset of patients, raising a need for response biomarkers [1]. A frequently used biomarker is the tumor mutational burden (TMB), a measure of the total number of mutations in the coding regions of the genome [1–2]. The U.S. Food and Drug Administration (FDA) has recently approved pembrolizumab, an immune checkpoint inhibitor (ICI) targeting PD1, for individuals with TMB-High (defined as ≥ 10 mutations/Mb, TMB-H) solid tumors [3]. Despite this approval, the effectiveness of TMB-H as a biomarker for stratifying responders to immunotherapy, termed here *TMB power*, differs considerably across cancer types [4,5], and the mechanisms underlying these differences have remained unknown. The tumor microenvironment (TME), including CD8+ T cells, dendritic cells, macrophages, B cells, T cell receptor (TCR) repertoire, and major histocompatibility (MHC) locus status, have been previously associated with the extent of immunotherapy response [6]. We thus hypothesized that differences in the immune activities in the TME of different cancer types may explain the variability observed in the *TMB power* across cancer types.

To study this hypothesis, we first collated the largest publicly available cohort of ICI-treated (anti-PD1/anti-PDL1) patient’s responses with TMB and demographic information, comprising 1959 patients [4, 5, 7, 8] together with an additional new cohort of 318 patients, comprising a total of 2277 patients across 14 cancer types. Analyzing this combined cohort, we first aimed to depict the association between TMB-H and patients’ response to ICI in each cancer type. To this end, we computed the difference in overall survival (OS) between patients with TMB-High vs TMB-Low, i.e., the Hazard ratio (HR) of survival, in each cancer type (**Figure 1A).** The HR is significantly < 1 for 8 out of 14 cancer types (using P<0.05 as a significance threshold), testifying to overall higher survival in patients with TMB-H, however, its magnitude varies considerably across cancer types. A similar trend and variability are observed for progression-free survival (PFS) **(Extended Figure 1)**. Repeating this analysis using the tumor response status and computing the odds ratio (OR) of objective response rate (ORR) between TMB-H vs low TMB patients, 5 out of 11 cancer types have a significant OR >1 (P<0.05, **Figure 1B**), testifying to a higher response rate in patients with TMB-H, but not in all cancer types. These observations are in line with previous findings [4,5], showing that the TMB-H biomarker based on a universal FDA-approved cut-off is predictive of response only in a subset of cancer types, with variable predictive power. We quantify the stratification power of TMB in identifying immunotherapy responders (termed *TMB power*) in each cancer type as 1/HR in terms of overall or progression-free survival and OR in terms of tumor response.

**Figure 1.**
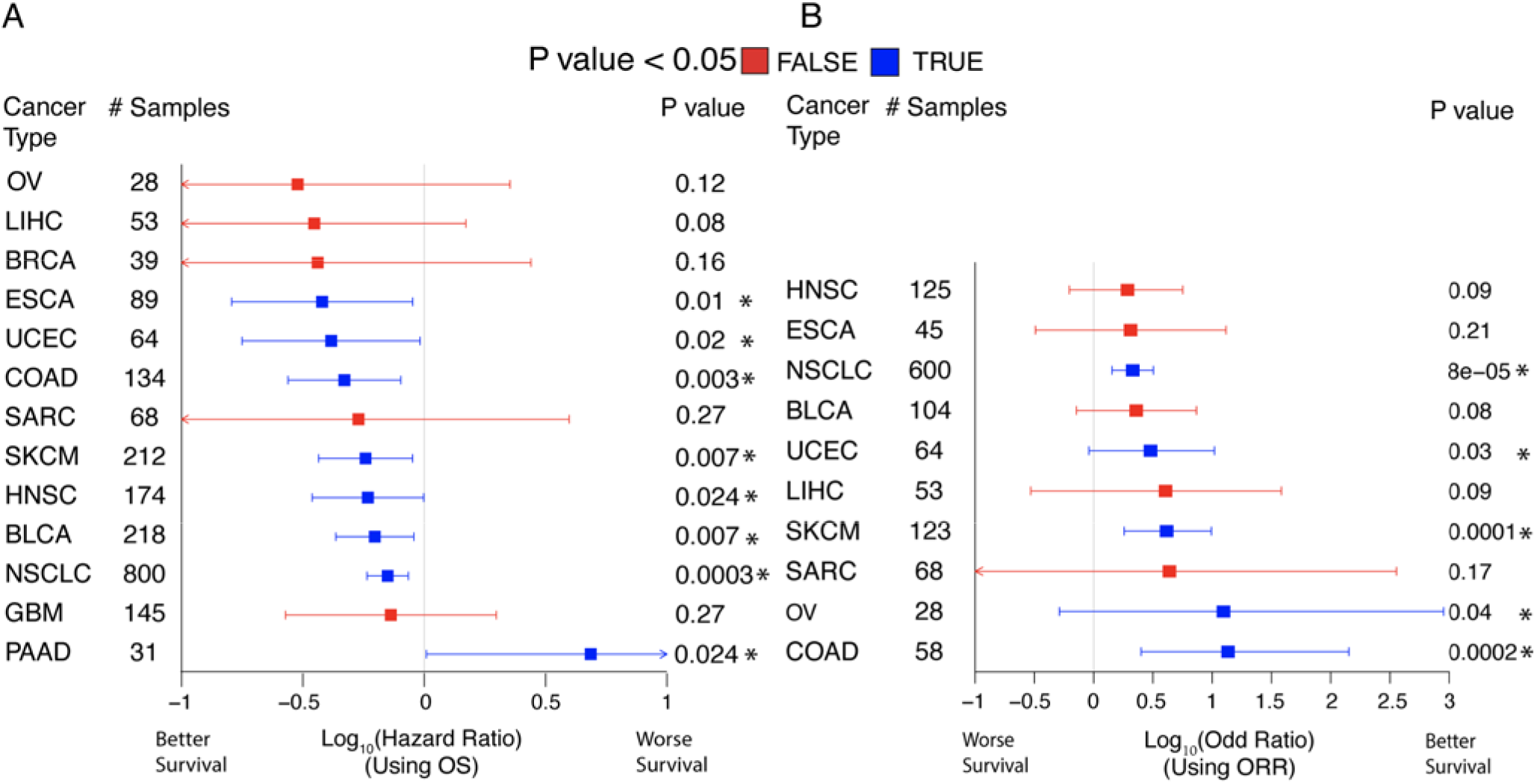
ICI response of TMB-H vs TMB low groups, for different cancer types. **(A)** Hazard Ratio of overall survival (OS) (x-axis) between patients with high vs low TMB computed using a Cox regression model. Cancer types having a significant hazard ratio are colored blue vs. red, denoting cancer types for which the HR is not significant. Error bars represent the 95% confidence interval and P-values were computed using a log-rank test. Cancer type abbreviations follow those used in the TCGA [9]. The number of patients with each cancer type is provided in the second column. **(B)** The results of a similar analysis using response status and the Odds Ratio. Renal cell carcinoma (KIRC) is not reported in (A) as its HR cannot be computed confidently.

We next quantified the mean levels of various immune-related factors in the TME of a given cancer type. To this end, we mined whole-exome sequencing and RNA sequencing data of pre-treated samples for the above 14 cancer types from The Cancer Genome Atlas (TCGA). In each cancer type, we estimated the mean levels of 31 different immune-related factors (**Extended Table 2**) that have been previously reported to be associated with ICI response [10]. Those include: (1) tumor neoantigen characteristics, including neoantigen hydrophobicity, intratumor heterogeneity, and neoantigen burden; (2) tumor microenvironment characteristics, including the abundance of different immune cells, the cytolytic score, T-cell exhaustion, and interferon-γ signatures, and T-cell receptor diversity, and finally, (3) checkpoint target–related variables, including PD-L1 protein expression, the combined positive score (defined as ratio of number of PD-L1 staining cells (tumor cells, lymphocytes, macrophages) out of the total number of viable cells) and fPD1 (the fraction of high PD1 staining tumors in a given cancer type).

To identify the immune-related modulators of *TMB power*, we computed the correlation between the mean-levels of each immune factor described above and the three measures of *TMB power* based on OS, ORR, and PFS, across the 14 cancer types we studied (**Figure 2A, B & Extended Figure 2A leftmost,** in respective order). Four immune factors emerge as being correlated with the *TMB power* for all three outcomes measures (**Figure 2C**). Two modulators are positively correlated with the *TMB power*, including *M1 macrophage levels* (correlation strength with TMB power based on OS is Spearman Rho = 0.61, P = 0.02, **Figure 2D-*Top-left panel***) and *tumor purity levels* (Spearman Rho = 0.44, P = 0.13, **Figure 2D-*Top-right panel***). We termed them *positive modulators*. Two other modulators are negatively correlated with the *TMB power* (*negative modulators*), including the *PDL1 combined positive score* (Spearman Rho = −0.39, P = 0.19, **Figure 2D-*bottom-left panel***) and *resting dendritic cells* (Spearman Rho = − 0.38, P = 0.21, **Figure 2D-*bottom-right panel***). Repeating the above analysis using only cancer types with statistically significant hazard ratios or odds ratios (**Methods**) yields concordant findings, where the same four modulators are the top ranked (**Extended Figure 2B-C**).

**Figure 2:**
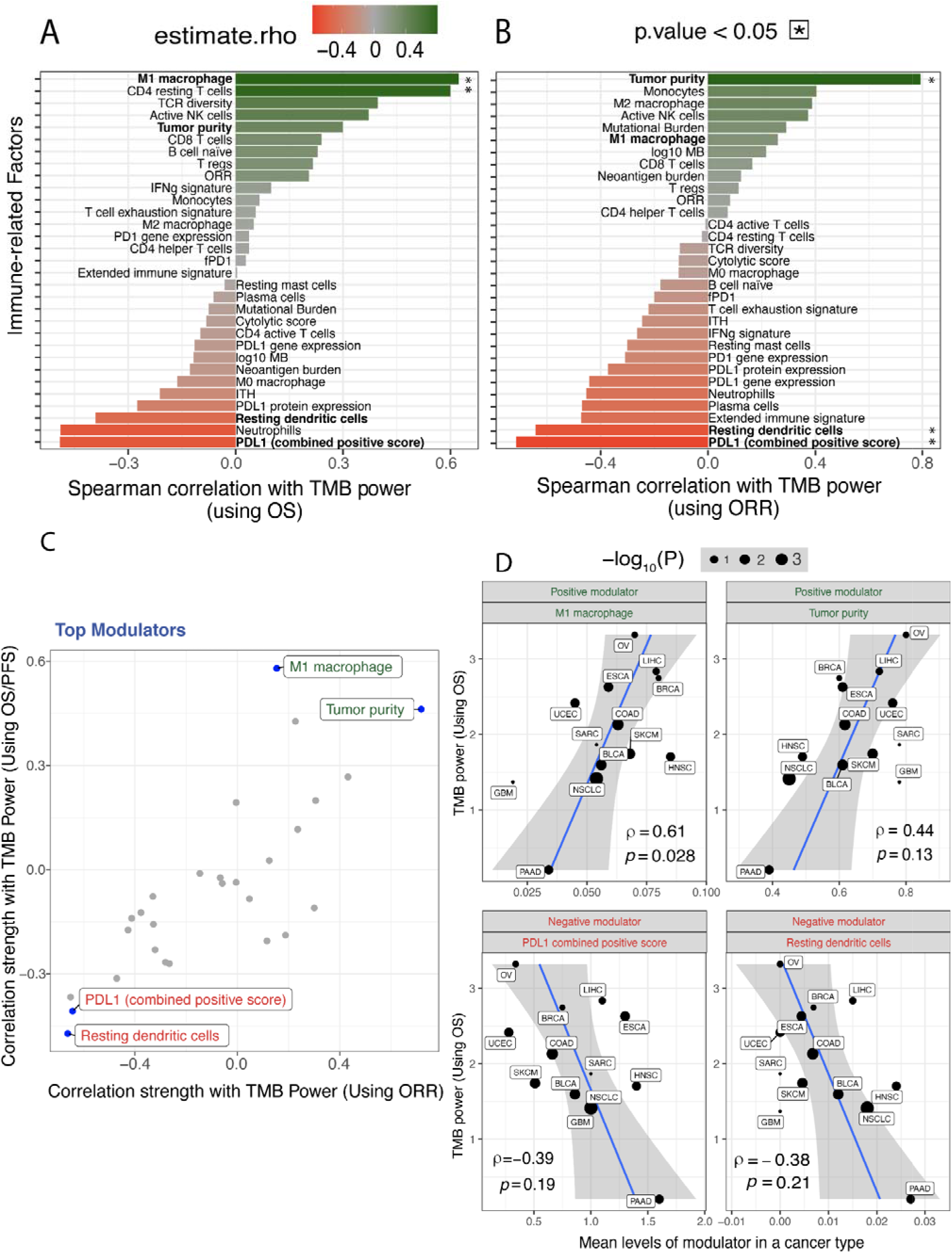
Immune modulators of *TMB power*: **(A)** Correlation strength between *TMB power* based on overall survival (OS) and levels of immune-related factors across 14 cancer types, computed using Spearman Rho (x-axis). Green and red bars denote positive and negative modulators, respectively, and the intensity of the color denotes the strength of correlation. Significant correlations (P<0.05) are marker with “*” and four highly correlated modulators are shown in bold. **(B)** This analysis is repeated to identify modulators of *TMB power* based on tumor response status (ORR). **(C)** The strength of correlation with TMB power using ORR (x-axis) and OS/ PFS (y-axis) is provided for each modulator, where the top four are highlighted and labeled. **(D)** Scatter plots showing the relationship between the mean levels of each of the four top modifiers in a cancer type (x-axis) and the TMB power (in terms of OS). The best fit line is provided in blue, where the shaded region denotes a 95% confidence interval. Renal cell carcinoma (KIRC) is not reported in **(D)** as its HR cannot be computed confidently.

We next built a multivariate linear model predicting *TMB power* at a given cancer type based on the levels of the four top modulators identified above, assessing their aggregate predictive power in a leave-one-out cross-validation procedure. We built this model separately for all three measures of *TMB power* (OS, PFS, ORR). The models performing best across all the three measures of *TMB power* only used two features - M1 macrophage and resting dendritic cells levels predicting *TMB power* based on OS with a Spearman Rho = 0.76, P = 0.0036 (**Figure 3A)**. Adding the two remaining modulators does significantly not improve model performance.

**Figure 3:**
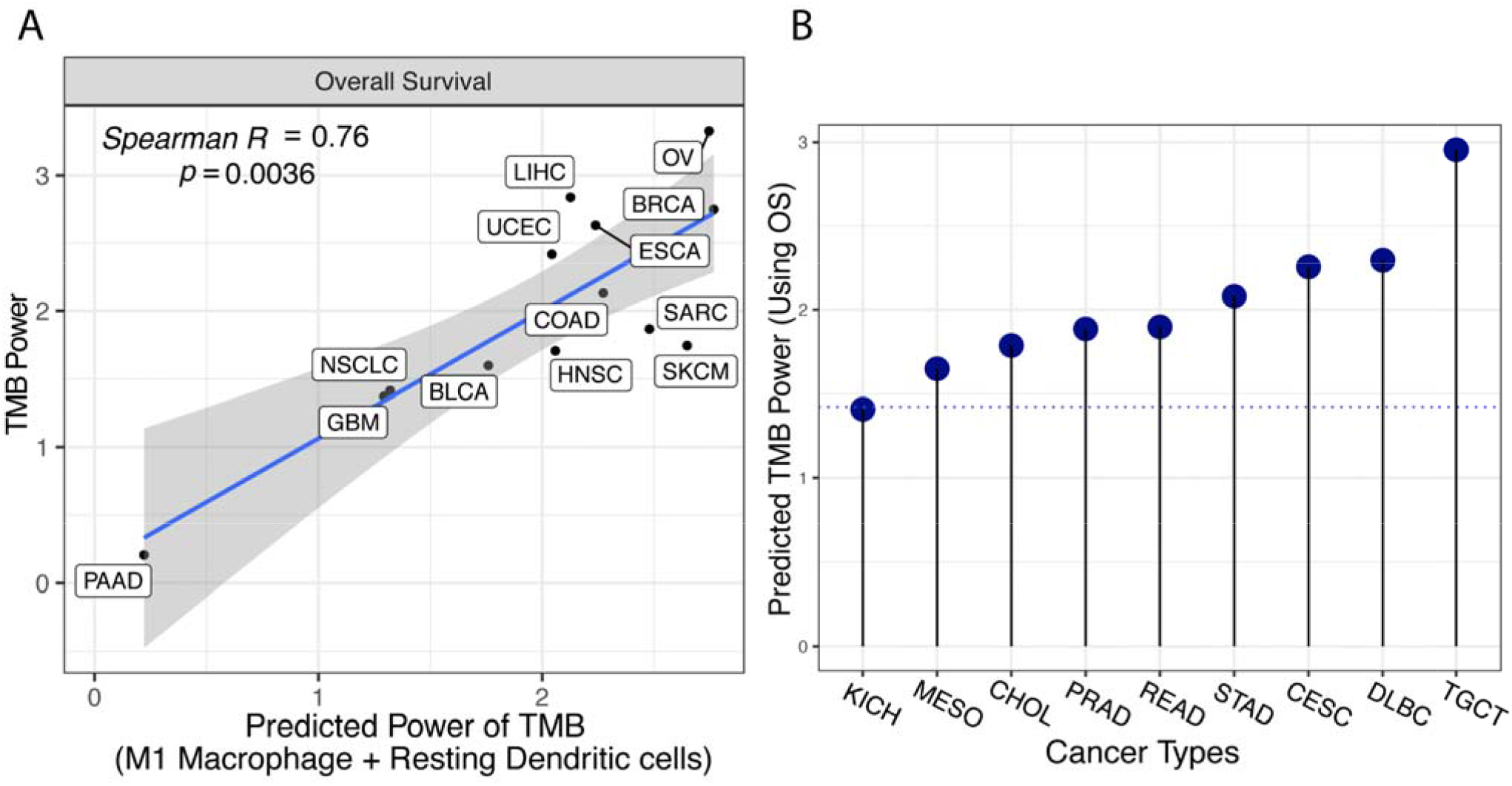
Predicting *TMB power* across cancer types. **(A)** The correlation between observed *TMB power* (y-axis) and its predicted value is, based on M1 macrophage and resting dendritic cell levels. The best fit line is provided in blue, where the shaded region denotes the 95% confidence interval. The spearman correlation and significance are provided at the left top corner. **(B)** Predicted TMB power using this model (y-axis) in 9 additional cancer types (x-axis), the blue dotted horizontal line shows the *TMB power* observed for NSCLC.

Using this two-feature linear model, we predicted the *TMB power* in 17 additional cancer types. These cancer types do not have publicly available TMB and ICI response data but their mean levels of M1 macrophage and resting dendritic cells could still of course be estimated from the TCGA cohort. We could confidently predict *TMB power* for 9 of these cancer types, where the two modulators’ levels were within the interpolation range of the regression (**Methods**). Notably, in 8 of these, the predicted *TMB power* was greater than that observed for lung cancer, where TMB-H patients have been shown to have a higher response rate and median survival in large clinical trials [11] (**Figure 3B, Extended Table 3)**. The top ranked cancer types are TGCT (Testicular Germ Cell Tumors, TMB power = 2.95, **two** times higher than the TMB power observed in lung cancer), DLBC (Lymphoid Neoplasm Diffuse Large B-cell Lymphoma, TMB power = 2.30, **1.6** times higher than the TMB power observed for lung cancer) and CESC (Cervical squamous cell carcinoma, TMB power = 2.26, **1.6** times higher than the TMB power observed for lung cancer). Among those, we think that cervical squamous cell carcinoma is probably the most interesting cancer type to further test the utility of TMB-H as a biomarker, as it has the highest overall immunotherapy response rate (20%, mined from [10]).

## Discussion

We identified two key immune-related factors whose levels are associated with the ability of TMB-H biomarker to stratify immunotherapy responders. Specifically, we find that high levels of M1 macrophages and low resting dendritic cells are predictive of cancer types with high *TMB power.* Aligned with these findings, M1 macrophages have been reported to provide an anti-tumor environment by fostering an inflammation response against tumor activating CD8 T cells, and thus their higher levels would likely augment the response to immunotherapy [12]. In contrast, resting dendritic cells provide a pro-tumor environment by inducing tolerance to tumor antigens via inducing T cell death or an anergic state (long-term inactivated state) or suboptimal priming, and thus their higher levels would likely suppress the response to immunotherapy [13, 14].

One limitation of our study is that our analysis is based on immune factors computed by deconvolution of bulk tumor data, which even though now being an accepted practice employed in many studies, only reflects estimations of different cell populations in the TME. Second, we should note that our data analysis combined data on patients receiving different formulations of anti-PD1 and anti-PDL1 for statistical power, whereas the FDA approval is for pembrolizumab (anti-PD1) specifically. Consequently, we expect that our results would be further refined as single cell based measurements of immune cells abundance and activity in different cancer types become available. Interestingly, we note that when we tested the predictive power of our modulators *at a patient-level* within a cancer type in four different cohorts [15–19], we did not find them to be predictive of TMB power (**Extended Figure 3**). Thus, as with TMB-H levels themselves, the factors determining response to immunotherapy across different cancer types are very different from those determining the response of individual patients within a given cancer type [20].

## Methods

### Data and preprocessing

All patients provided informed consent to an MSK institutional review board-approved protocol, permitting the return of results from sequencing analyses for research. We collated the 1. A publicly available cohort of ICI treated (anti-PD1/PDL1) patient’s responses with TMB and demographic information, comprising 1959 patients, and 2. data from an additional 318 patients, yielding a total 2277 patients across 14 cancer types. We removed the cancer types where all the patients have TMB < 10 mut/MB.

### Computing TMB power for each cancer type

We first compute the hazard ratio (HR) of survival (overall and progression-free) and odds ratio (OR) of response rate between TMB-H vs low group. In case of overall and progression-free survival, the TMB power is 1/HR. And, in case of tumor response status, TMB power is defined as OR.

### Mean levels of Immune factors across samples of a cancer type

These levels were mined from our previous publication [10] where a detailed methodology is provided. Please refer to the method section of [10]. We remove three immune factors - “active dendritic cells”, “resting NK cells”, and “active mast cell” from our initial immune factors set, due to low variance (0 in more than 50% of cancer types).

### Finding the immuno-modulators of TMB power

The correlation between TMB power and mean-level of 34 immune-related factors was calculated using Spearman rank correlation (rho). From this analysis, four candidate modulators consistently strongly correlated with TMB power across 1. different measures of outcomes, 2. for both cancer types sets, A. all and B. only ones with significant HR, 3. Both Pearson and Spearman correlation methods are selected for further analysis.

### Multivariate regression model to predict *TMB power*

We next built a multivariate linear regression model based on the above four modulators, using a standard leave-one-out cross-validation method. Here, we built a model using all possible combinations of features exhaustively and the performance of the prediction was evaluated based on Spearman rank correlation (rho). Using the best model, we next predicted *TMB power* for 17 additional cancer types where it is unknown. Among these, we noted that feature values are in the interpolation range i.e. range which our model was trained on, for only 9/17 cancer types and thus we restricted our prediction for these cancer types.

## Data and Code availability Statement

The study’s scripts and data are provided to reproduce each step of results and plots in <u>this</u> GitHub repository.

## Acknowledgments

This research used the computational resources of the NIH HPC Biowulf cluster (http://hpc.nih.gov). This research was supported by the Intramural Research Program of the National Institutes of Health, NCI, by Fundación Alfonso Martín Escudero (to CV) and the National Institutes of Health Cancer Center Support Grant P30 CA008748. S.S is supported by the NCI-UMD Partnership for Integrative Cancer Research Program. L.M is supported by the NIH R01 DE027738 grant.

## Author contributions

ER and SS conceived and supervised the study. NS and SS designed and developed the methodology. NS, SS, and CV acquired and analyzed the data. AAS, TAC, and LGTM advised on the methods and interpretation of results. SS, NS, and ER wrote the manuscript. All authors carried out a critical revision of the manuscript for important intellectual content.

