## Supplementary figures for "Immune determinants of the association between tumor mutational burden and immunotherapy response across cancer types"

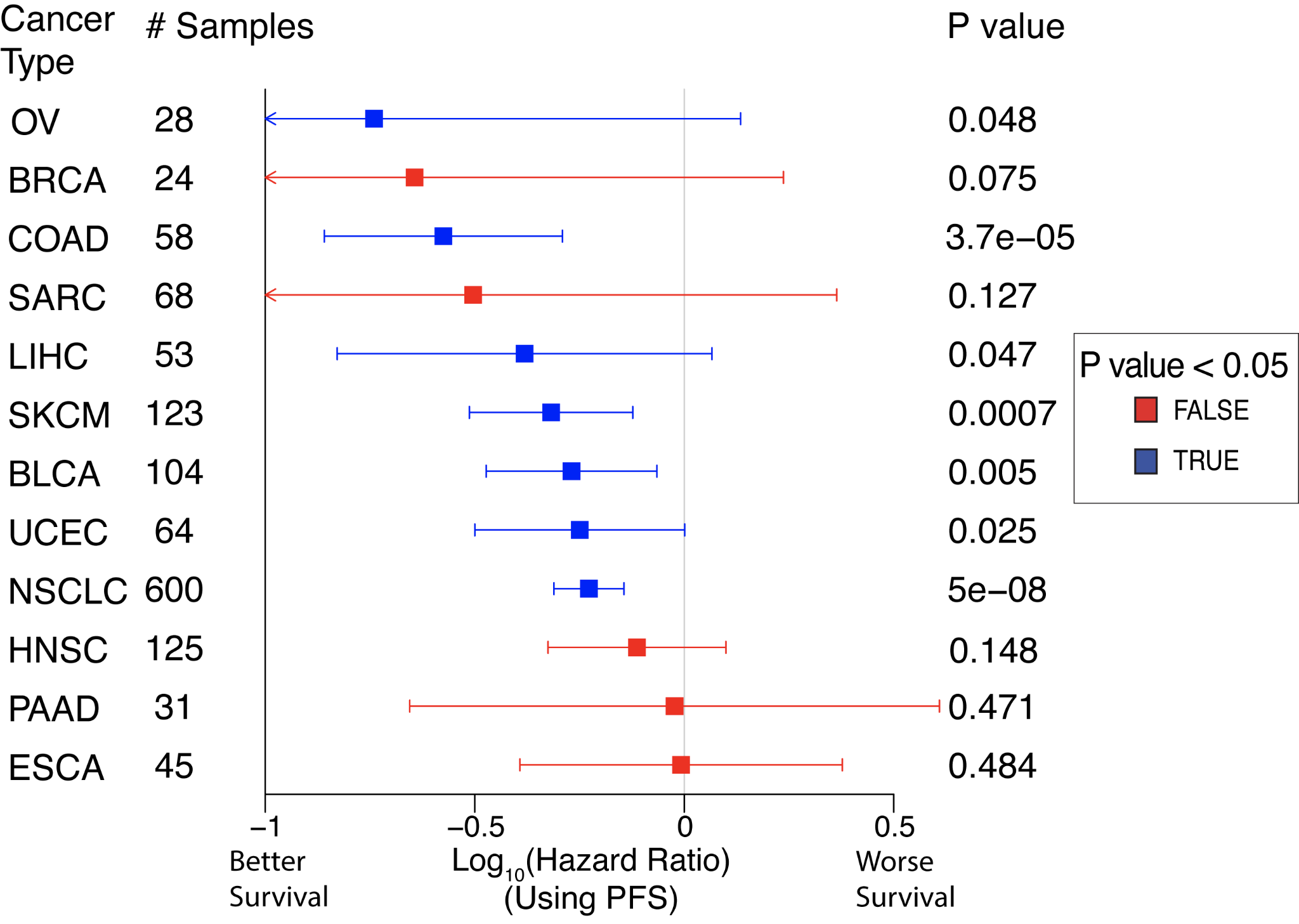

**Extended Figure 1: Hazard Ratio of progression-free survival after ICI treatment between TMB-H vs. TMB low groups. (A)** Hazard Ratio of survival (x-axis) between patients with TMB-H vs. TMB low computed using a Cox regression model, where the cancer types with significant vs. insignificant hazard ratios are color coded (blue vs. red, respectively) and the legend is provided at the right of the figure. Here, the error bar represents the 95% confidence interval and the significance is computed using a log-rank test. The cancer type abbreviation is used as in TCGA [9]. The number of patients available for the analysis in each cancer type is provided in the second column.

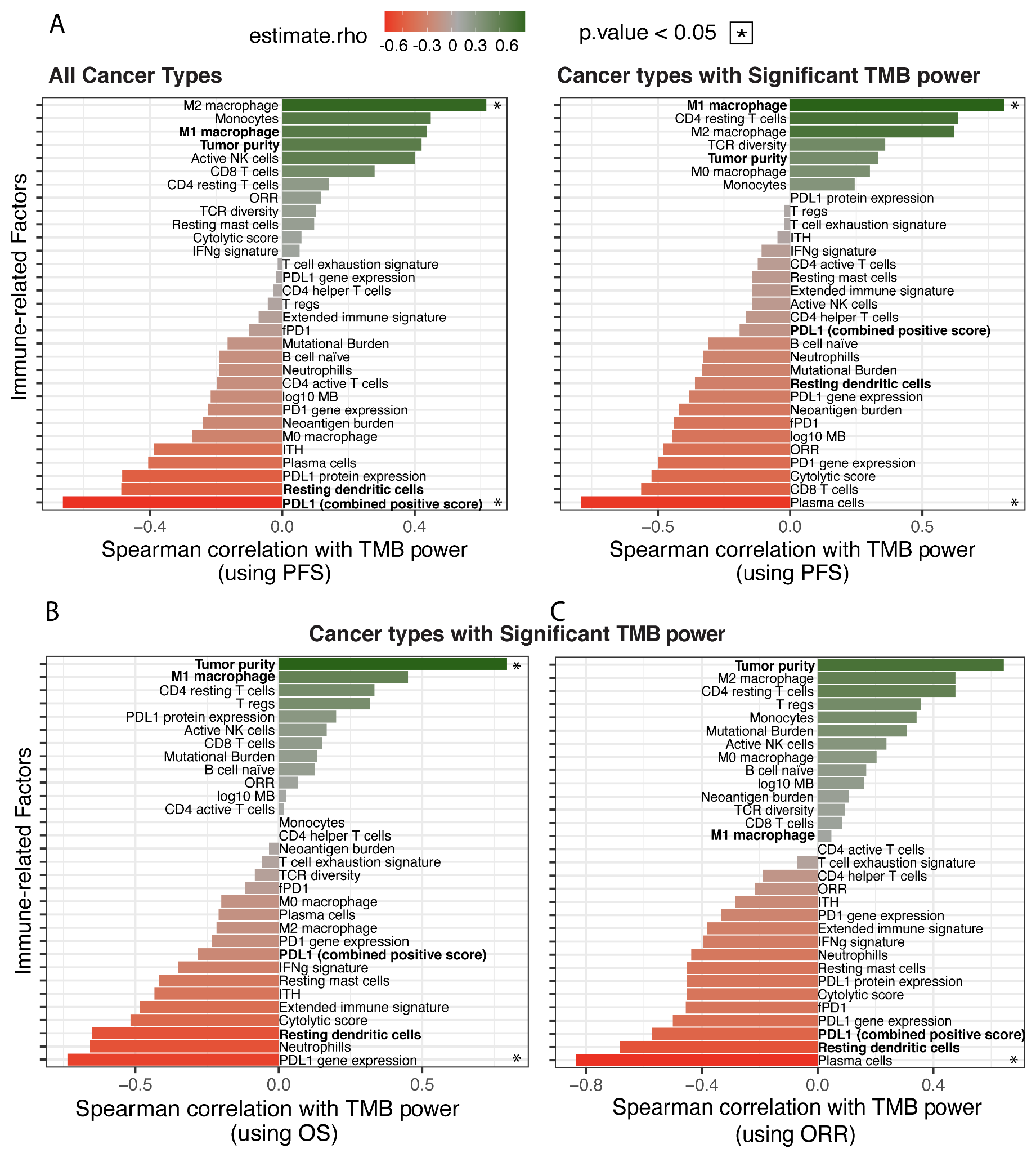

**Extended Figure 2: Statistical relationships between TMB power and three measures of response. (A)** Correlations between *TMB power* based on progression-free survival and mean levels of immune-related factors across 14 cancer types computed using Spearman Rho (x-axis), where the name of each factor is provided right beside the bar. The bars are color coded based on the legend on the top. This is repeated using *TMB power* based on progression-free survival (provided in **A rightmost**), overall survival (provided in **B**) response status **(**provided in **C)** across only cancer types with significant HR values.

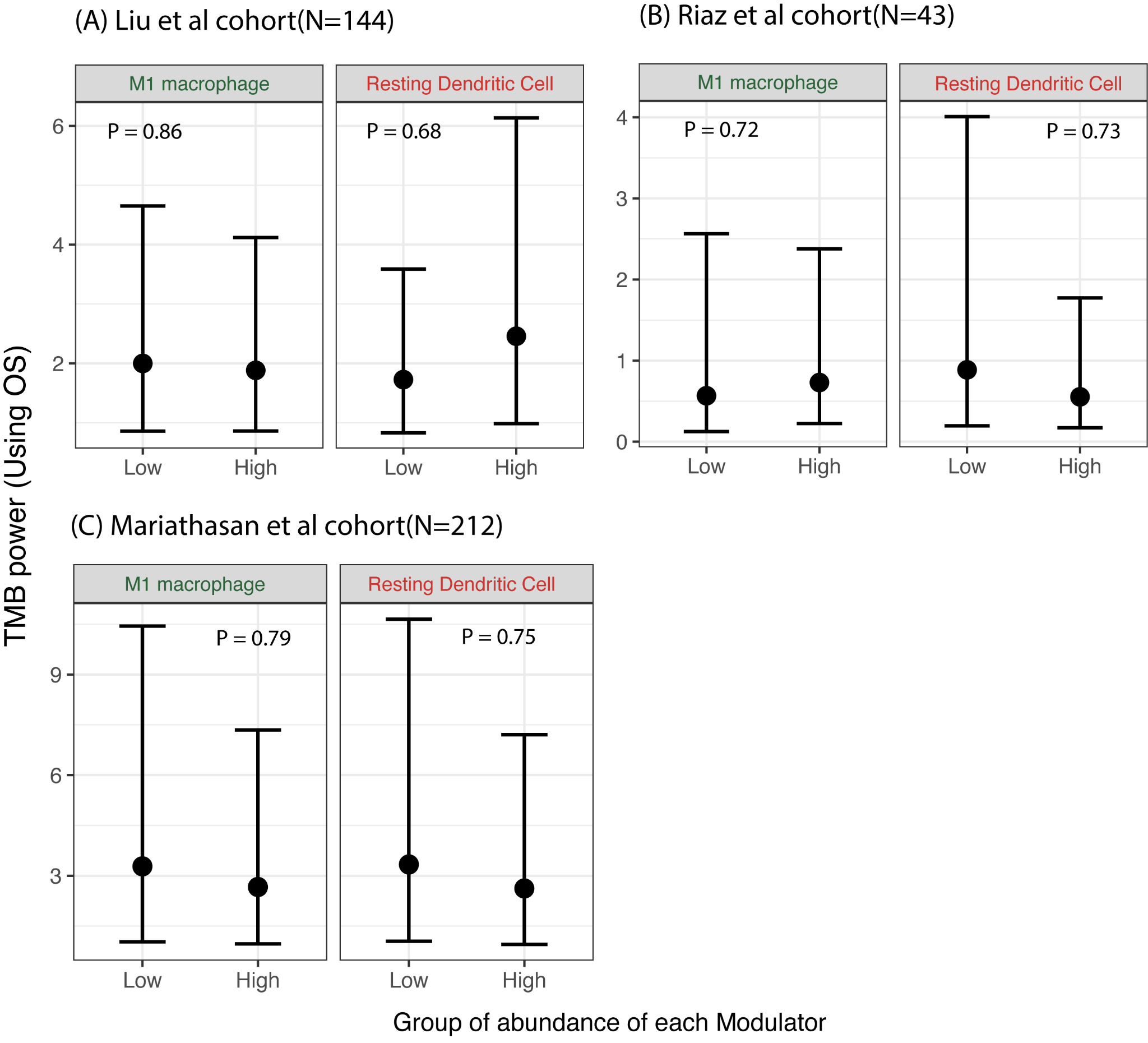

**Extended Figure 3:** Difference in *TMB power* between TMB-H vs TMB low, when cohorts are stratified by levels of two major modulators - M1 macrophage and resting dendritic cell, in three **cohorts.** *TMB power* is not shown for Hugo et al as the Cox regression did not converge likely due to the low number of samples.
